# Extended regions of suspected mis-assembly in the rat reference genome

**DOI:** 10.1101/332932

**Authors:** Shweta Ramdas, Ayse Bilge Ozel, Mary K. Treutelaar, Katie Holl, Myrna Mandel, Leah Solberg Woods, Jun Z. Li

## Abstract

We performed whole-genome sequencing for eight inbred rat strains commonly used in genetic mapping studies. They are the founders of the NIH heterogeneous stock (HS) outbred colony. We provide their sequences and variant calls to the rat genomics community. When analyzing the variant calls we identified regions with unusually high levels of heterozygosity. These regions are consistent across the eight inbred strains, including Brown Norway, which is the basis of the rat reference genome. These regions show higher read depths than other regions in the genome and contain higher rates of apparent tri-allelic variant sites. The evidence suggests that these regions may correspond to duplicated segments that were incorrectly overlaid as a single segment in the reference genome. We provide masks for these regions of suspected mis-assembly as a resource for the community to flag potentially false interpretations of mapping or functional results.

## Background and Summary

The laboratory rat (*Rattus norvegicus*) is an important model organism for studying the genetic and functional basis of physiological traits. Compared to the mouse, the rat shows a greater similarity to humans in many complex traits (Iannaccone and Jacob 2009) and has been widely used in physiological, behavioral and pharmacological research. With the arrival of high-throughput genotyping and sequencing technologies, the rat has also been used in genetic studies to map causal loci, or identify genes that affect disease-related traits.

An essential resource in such genetic studies is the rat reference genome (Gibbs et al. 2004), which provides the coordinate system to orderly manage our rapidly increasing knowledge of rat genes, their regulatory elements, gene products and variants, functional profiles of diverse tissues, as well as biological dysregulation in disease models. The reference genome is also the basic map in comparative analyses that focus on the evolutionary relationship among rat strains or between the rat and other organisms.

Gene discovery studies using animal models can be roughly classified by the type of mapping populations adopted. Currently, popular genetic systems involving the rat include naturally occurring outbred populations, laboratory-maintained diversity outbred populations, inbred line-based crosses (e.g., F2-crosses, or advance inbred lines), recombinant inbred lines, and many others. Regardless of the system, a comprehensive knowledge of DNA variation in the mapping population is essential for both the study design and biological interpretation. In this study, we sought to use whole-genome sequencing (WGS) to ascertain DNA variants in eight inbred strains: ACI, BN, BUF, F344, M520, MR, WKY and WN, which are founders of the NIH Heterogeneous Stock (HS) population. The HS rat has been used in genetic studies of metabolic and behavioral traits (Solberg Woods et al. 2010a; Solberg Woods et al. 2010b; Solberg Woods et al. 2012; Woods and Mott 2017; Holl et al. 2018; Keele et al. 2018). WGS data for these eight strains have been previously described (Baud et al. 2014; Hermsen et al. 2015), using the SOLiD technology. Here we present WGS results from the Illumina technology, containing genotypes at ~16.4 million single-nucleotide variant (SNV) sites. We expect that the sequences of the eight HS founders and fully-ascertained DNA variations can aid the imputation, haplotyping, and fine mapping efforts by the rat genomics community.

When analyzing the SNV data we noted that, while the eight founders are inbred, all contain an unusually high amount of heterozygous nucleotide positions. Remarkably, these sites tend to concentrate in hundreds of discrete genomic regions, which collectively span 6-9% of the genome. We show that the heterozygous genotypes tend to recur in multiple, if not all, of the eight strains, and that the suspected regions tend to have higher-than-average read depths. We propose that these regions can be explained by mis-assembly of the rat reference genome, where many of the highly repetitive segments may exist in tandem or dispersed in distant regions, but have been erroneously "folded" in the current coordinate system, causing the reads of high homology that originate in distinct homozygotic regions to falsely aggregate to produce apparent heterozygous calls. This interpretation is not unexpected when one compares the genome assembly statistics between mouse and rat: the latest release of the mouse reference genome, GRCm38.p6, contains 885 contigs and a median contig length of 32.3 megabases; whereas the rat reference genome, rn6, has 75,687 contigs and a median length of 100.5 kilobases. With this report we release mask files for the suspected regions, so that they can be used to flag questionably results in current genomic studies until the time when a revised, more accurate reference assembly becomes available.

## Methods

### Animals, DNA samples, whole-genome sequencing

Eight animals, one for each of the eight founders of the HS population, were used in the study. Genomic DNA of the original founder strains was obtained from NIH (M.M.). DNA was extracted at the University of Michigan (Ann Arbor, MI) from the liver of seven animals (ACI/N, BUF/N, F344/N, M520/N, MR/N, WKY/N, WN/N), and from the tail of a BN/N animal. All animals were female. The DNeasy Blood and Tissue Kit from Qiagen (Hilden, Germany) was used for DNA extraction. Samples were further QC’d and sequenced at *Novogene* (Beijing, China) following the standard Illumina protocols. Library preparation produced fragment libraries of ~350 bp insert length. Sequencing was done on Illumina HiSeqX-Ten to collect 150 bp paired-end data, aiming for an average depth of 25X per strain.

### Sequence alignment and variant calling

We aligned the raw sequence reads to the rat reference genome (rn6) using *BWA* version 0.5.9 (Li and Durbin 2009), removed duplicates using *Picard* v1.76 (Ramdas 2018), and performed realignment, recalibration and joint variant calling across eight strains with the UnifiedGenotyper with *GATK* v3.4 (McKenna et al. 2010). We removed variant sites with fewer than 10 reads in eight samples, and variant site quality score (QUAL) <= 30. We chose not to use the HaplotypeCaller as we have only eight inbred lines, which are not the population-based samples suitable for building haplotypes.

ChrX data showed the same pattern of heterozygosity as the autosomes in all eight animals, thus confirmed that they are female. We excluded the Y chromosome calls in downstream analysis. We did not call indels in this data release.

For comparison purposes we also ran the analysis with (1) two earlier versions of the reference genome, rn4 and rn5; (2) two other aligners. The first is by feeding the *BWA* aligned files into *Stampy* (Lunter and Goodson 2011) version 1.0.32. *Stampy* alignment shows higher sensitivity than BWA, especially when reads include sequence variation (Lunter and Goodson 2011). (The use of BWA-alignment as input for Stampy is to increases alignment speed without reducing sensitivity). The second is *Bowtie2* v2.1.0 (Langmead et al. 2009). All the post-alignment processes followed the same *Picard* and *GATK* steps. The results show high concordance among the genome versions (**Supplementary Table 1**) and among the aligners (**Supplementary Table 2, 3**).

### Comparison with the previously published variant calls using SOLiD

We compared our variant call set (for *BWA* alignment and rn6) with that by Hermsen et al (2015), which was based on the SOLiD sequencing data. As that call set was aligned to rn5, we lifted over the variants to the rn6.

### Defining regions of unusually high rates of heterozygosity

The final call set contains >16.4M SNV sites. We divided the genome into 1000-site windows, with a median window length of 221,100 bases. For each of the eight samples and in each window we computed the fraction of heterozygous sites (only using the number of non-missing sites in that window as the denominator). Based on the inflection point in the empirical distribution of this per-window heterozygosity fraction (**Supplementary Figure S1**) we chose a cutoff of 25% to designate windows as of high-heterozygosity. We concatenated adjacent windows of high-heterozygosity into the same segment, and in a second step, merged adjacent high-het segments if they are separated by a single “low-heterozygosity” window, if that window had more than 0.175 heterozygote rate (**Supplemental Figure S5**). After merging, there is no evidence of many very short low-het segments separating high-het segments (**Supplemental Figure S2**).

## Data Records

We share data files to cover multiple levels of processing.

1. Raw FASTQ/BAM files will be deposited in the *Rat Genome Database* (RGD) (Shimoyama et al. 2015).
2. Variant calls from the UnifiedGenotyper (using *BWA* and rn6), as VCF files for the eight strains, on our GitHub page (Ramdas 2018).
3. Mask files, also on our GitHub page, documenting the regions of high heterozygosity, for each of the eight founder strains, as well as regions that are highly-heterozygous in all the eight.

## Results

### Description of the variant calls

DNA from one female animal for each of the eight strains was sequenced **(Methods)**. Median read depth over the genome ranges 24X - 28X across the eight samples. Joint variant calling revealed 16,405,184 post-filter single-nucleotide variant sites on the autosomes and chromosome X. The number of heterozygous sites per strain varies from 1,560,708 (BN) to 2,114,990 (WN) **(Supplemental Table 4)**. BN represents the reference genome, and has more Ref/Ref than Alt/Alt genotypes. In contrast, the other seven strains have a comparable number of Ref/Ref calls as Alt/Alt calls.

We compared our genotype calls with those from Hermsen et al. (2015) by calculating between-study concordance rates at sites reported in both, and using genotypes that do not include the missing calls. **Supplemental Table 5** shows that each of the eight lines can be correctly matched between the two datasets, confirming the sample identity even when the two studies were based on different animals for a given line. BN has the highest between-study concordance: 0.95. Six other lines have concordance > 0.86. However, MR has the lowest concordance, 0.69. To determine if an animal from another line was mislabeled as MR, we compared our MR data with a larger panel of 42 strains previously published (Hermsen et al. 2015) and found that our MR had the highest match with MR and WAG-Rij in that study. This suggests that the MR lines in different laboratories may have diverged to an unusually large degree.

### Regions of unexpected high-heterozygosity

The eight founder strains were kept as inbred lines over many generations, thus were expected to show low heterozygosity across the genome (Brenner et al. 2002). However, when we calculated the fraction of heterozygous genotypes in consecutive 1000-SNV windows, we observed highly varied distribution of this metric along the genome. Not only were there many windows of high heterozygosity (>0.25), they also tended to recur in multiple lines **(Figure 1)**. Some of these windows were found in all 8 strains, including BN, the strain of the reference genome **(Supplemental Figure S3)**.

**Figure 1.**
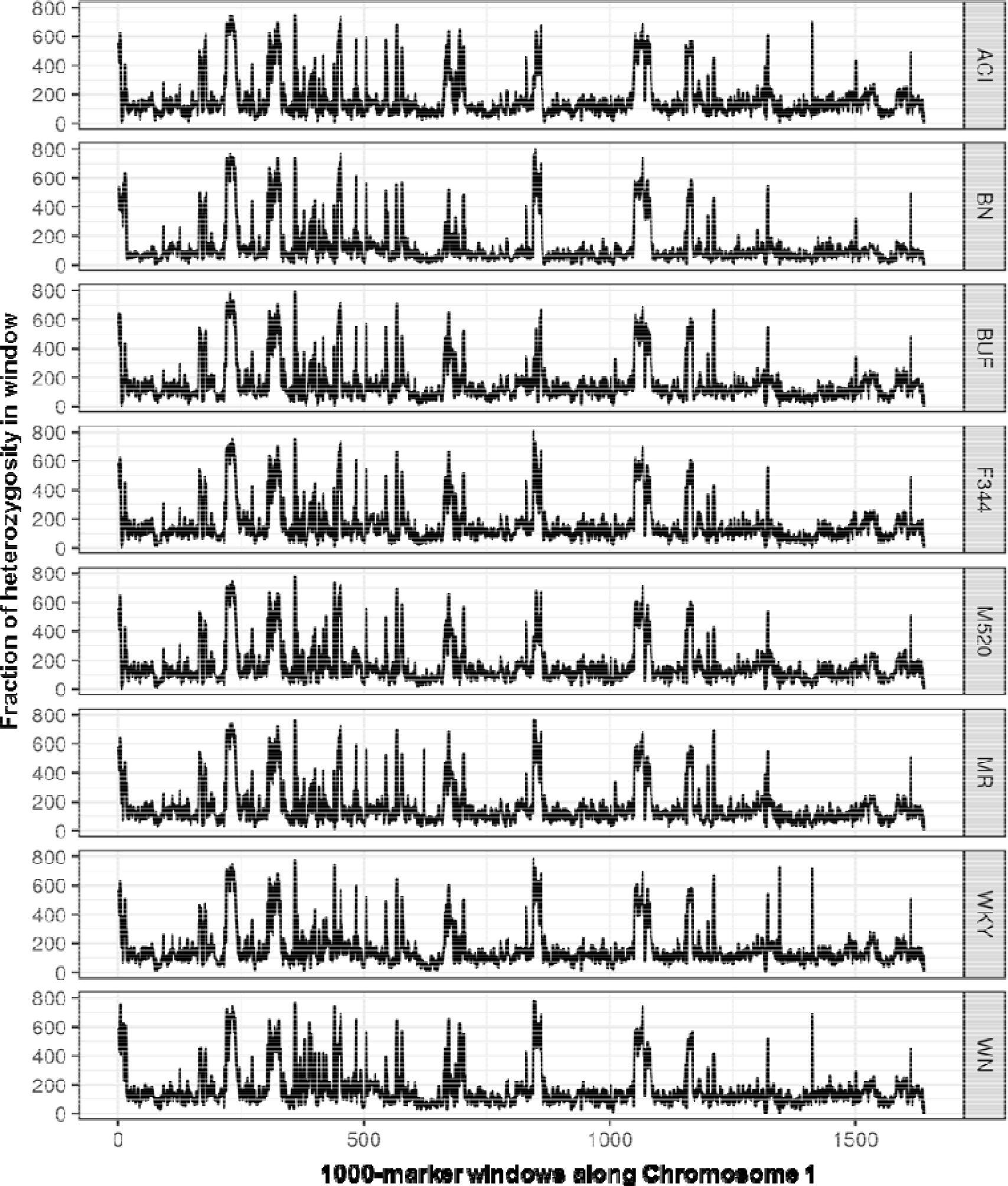
Consistent heterozygosity patterns across the eight lines. Shown are the fractions of heterozygous genotypes (y-axis) in non-overlapping 1000-SNV windows in chromosome 1, displayed for the eight inbred lines.

We chose a het cutoff value of 25% based on the distribution of per-window heterozygosity **(Supplemental Figure S1)**. After merging neighboring high-het windows if they are separated by a single low-het window with het rate > 0.175, we obtained 304-482 contiguous high-het regions across the eight lines (median 452), covering 176.4-254.8 Mb, equivalent to 6.3%-9.2% of the genome (average 8.4%). The distribution of segment lengths is shown in **Supplemental Figure S2**. Of the heterozygous calls in each line, 28-31% fall in the high-het regions (mean 29%). Of the 534,266 triallelic variant sites, 484,583 (~91%) fall in the regions that are high-het in at least one strains, covering 12% of the genome.

The high-heterozygous regions found in all eight lines contain 1,756 Ensembl genes. These genes were enriched for G-protein coupled receptors and olfactory receptors (fold enrichment of 3.2 and 2.0 respectively, Benjamini-Hochberg FDR < 0.01), which are known to have many paralogous copies in mammalian genomes (Hughes *et al*. 2018). There are 4,963 missense variants, 123 stop-gain variants, and 154 splice donor/acceptor variants in 372 genes in these regions.

### Heterozygous calls tend to be recurrent and show higher read depths

While **Supplemental Figure S3** shows that 1000-variant windows of high heterozygosity tend to appear in multiple lines, we also analyzed individual heterozygous genotypes to see how much they tend to recur in multiple lines. We divided the ~16.4M variant sites based on the number of heterozygous genotypes observed in the eight lines, thus defining nine variant site categories, from 0 to 8 heterozygotes **(Figure 2)**. If the heterozygous genotypes appear independently in the eight lines with a probability of p, the expected chance of seeing a site with two heterozygotes (that is, in two of the eight lines) would be proportional to p^2^, and in three lines: p^3^. We estimated the upper limit of p by counting the fraction of heterogeneous genotypes over all 8 lines in all sites, knowing that this fraction is already biased upward due to the highly recurrent heterozygous sites. The expected probability of seeing k heterozygous sites, a(k)p^k^, where a(k) is the sampling coefficient of the binomial distribution, drops much faster than the observed k-het counts **(Figure 2)**. For instance, the observed number of sites that are heterozygous in all eight lines is more than five orders of magnitude higher than expectation.

**Figure 2.**
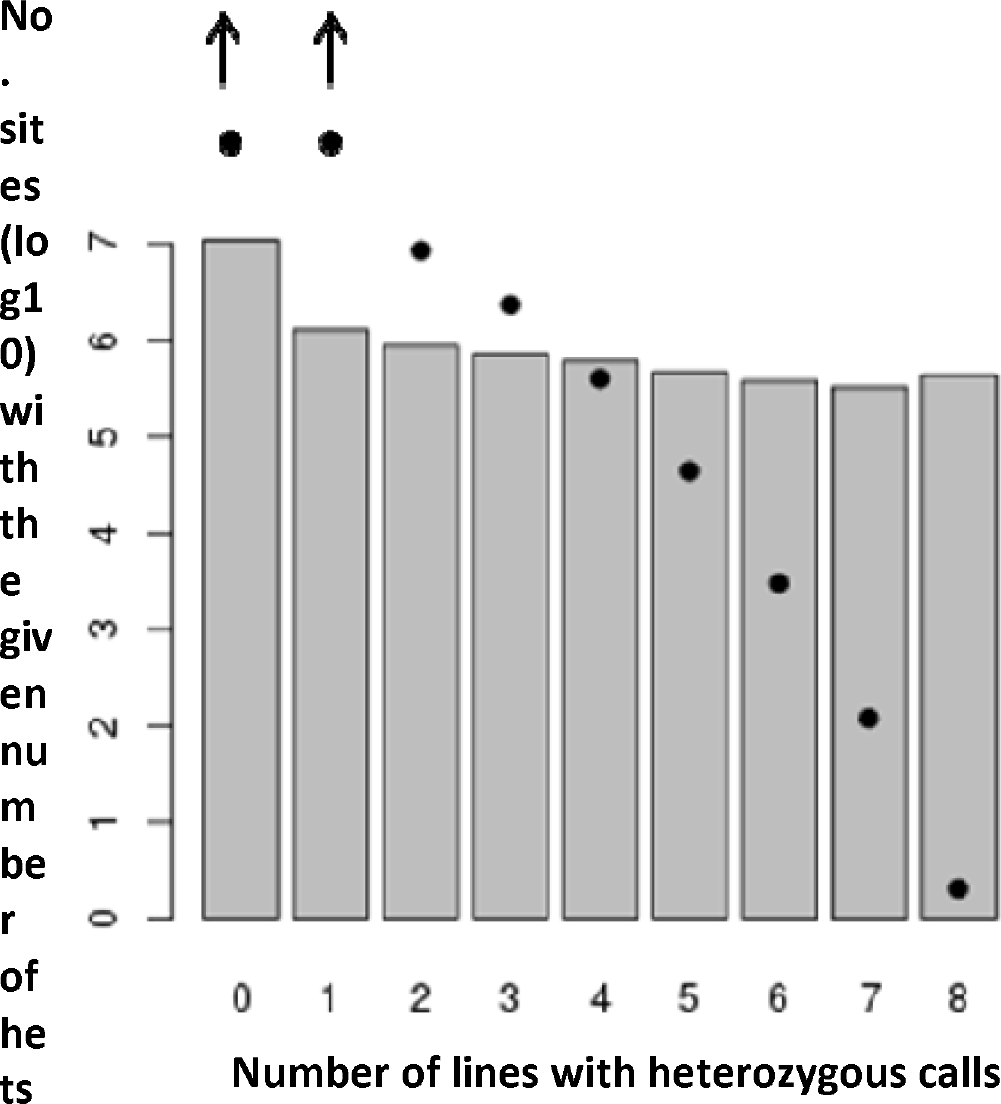
Recurrence pattern of heterozygous genotypes across 8 founders. All variant sites were categorized as having 0, 1,…, up to 8 heterozygous genotypes. The bar graph displays the histogram of the observed sites, while the dots show the expected number of sites if recurrence is random, estimated under a simple binomial model. The expected values for 0 and 1 founders (7.21 and 7.20 respectively) exceed the limits of the plot.

The tendency for an individual site to appear heterozygous in multiple lines is related to higher read depth at these sites **(Figure 3A). Figure 3B** shows an example where low-het regions tend to have lower, and more stable, read depth than nearby regions of high heterozygosity. Notably, the read depth is higher in high-het regions for both homozygous and heterozygous genotypes.

**Figure 3.**
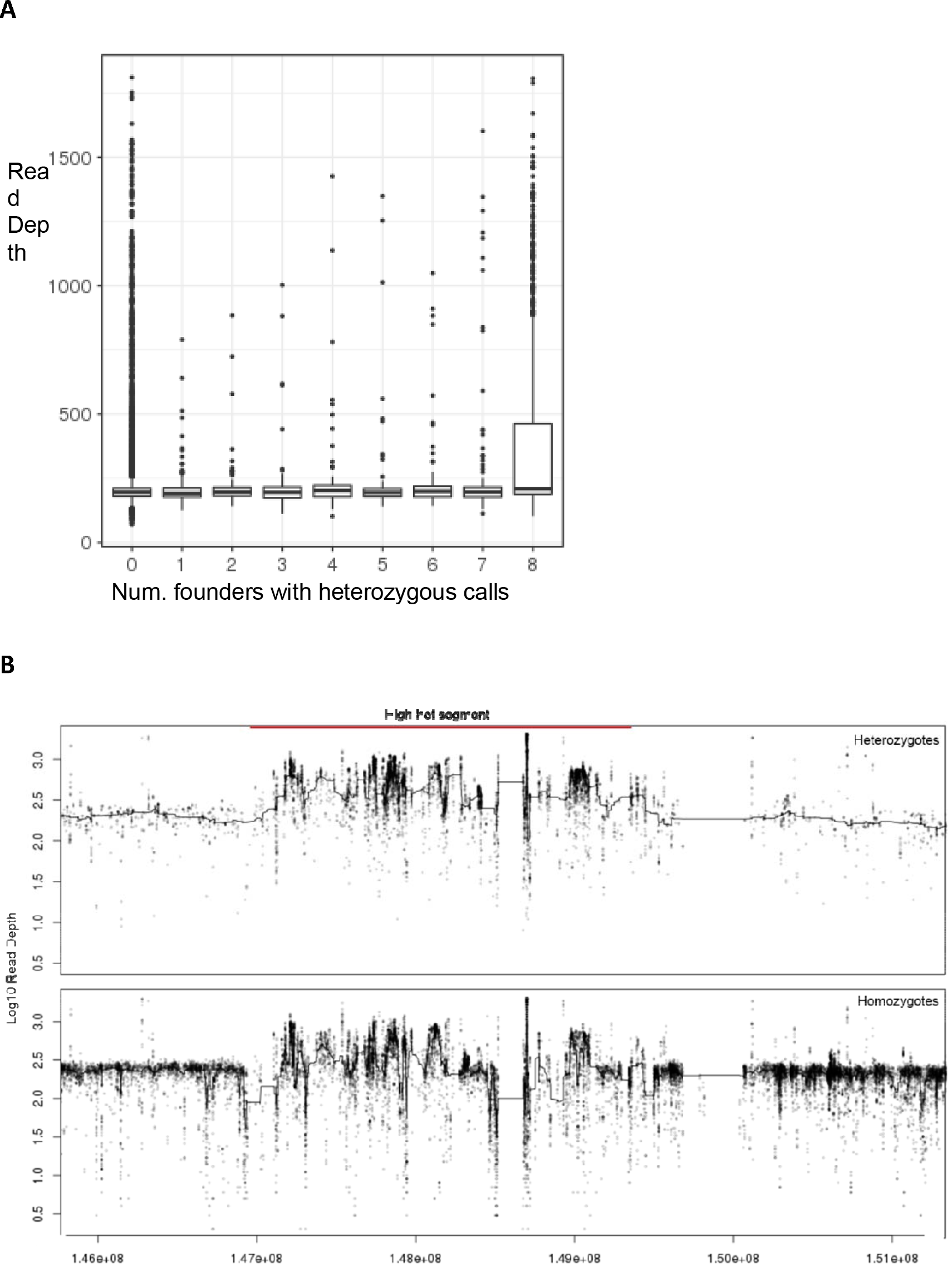
Higher read depths in high-heterozygosity windows. **A**. Boxplot of per-window average read depth stratified on the x-axis by the number of lines for which a given window is classified as high-het. It shows that windows with high-het in all eight lines tend to show higher average read depth. **B**. An example of a 5Mb region in Chr1 and line AC, with a high-het region in the middle. Y axis shows the read depth for individual sites, showing that in the high-het window, both the het calls (upper panel) and hom calls (lower panel) show higher read depth and often higher variance of read depth.

### Concordance with results from a different platform

We analyzed the previously released variant calls from Hermsen *et al*. 2015 (after lifting over these calls to the rn6 genome from the original rn5) to see if our high-heterozygosity segments were also found in their calls. The Hermsen dataset revealed 5-18% (median 14%) of the genome in high-het windows. We found that these high-het regions tend to match in the two call sets (an example in **Supplemental Figure S4)**, with an overlap of 24%-36% as defined by the length of the intersect of segments called high-heterozygosity in both calls, divided by the union of regions defined as high-heterozygosity in either call.

We have compiled a list of these high-het regions of the genome and provided them as bed files on our Github page: <u>http://github.com/shwetaramdas/maskfiles/</u>.

## Discussion

In this study we created WGS-based variant call sets for eight inbred lines which represent the genomic source of the multi-parental HS population. Our data suggest that the current rat reference genome contains hundreds of problematic regions where inbred lines show increased apparent heterogeneity and higher-than-usual read depth. These regions make up ~8.4% of the genome, and likely represent regions of mis-assembly. In other words, a properly unfolded genome may be ~8% longer.

Several lines of evidence led us to the interpretation that regions of high-heterozygosity likely represent an artifact where two or more segments of high similarity are incorrectly "collapsed", or "folded", into the same segment in the rat reference genome. First, the rate of heterozygous calls is higher than what one would expect for inbred lines; and these heterozygous calls cluster in discrete regions. Second, these high-heterozygosity regions tend to recur, sometimes in all eight lines, indicating that they are unlikely to have arisen from genetic drift or strain-specific selection. Finally, these regions tend to have higher read depth than surrounding regions. The rat reference genome, unlike the human and the mouse reference genomes, was assembled using a hybrid of shotgun-sequencing and clone-based approaches. Our results suggest that this mixed (primarily shotgun) approach has not been sufficient to resolve all the repetitive regions. Thus, Illumina sequence reads that originated in distinct but highly similar regions–some of these may fit the strict definition of paralogous regions–are incorrectly aligned to the same collapsed region in the reference genome, producing false heterozygous calls.

Guryev *et al* (Guryev et al. 2008) studied the rat reference genome (rn4 at the time) and used the read-depth distribution to identify 73 regions of suspected mis-assembly, which make up ~1% of the genome. Only 2 of these 73 regions can be lifted over to the rn6 genome and both were called high-het in our data. However, they didn’t analyze the distribution of heterozygous genotypes. Another study performed WGS on the inbred strain SHR/Olalpcv and observed higher-than-expected levels of heterozygosity. The authors suggested that this could have resulted from collapsing of reads from segmental duplications (Atanur et al. 2010), an interpretation similar to ours. Because they analyzed a single strain, the data did not show the consistency of suspected regions across strains.

In an early draft of the mouse genome segmental duplications were inadvertently misassembled (Bailey et al. 2004). Our estimates of misassembled regions underscore a similar problem with the current rat reference genome. These regions harbor more than 1,700 genes, with 5,000 apparent missense variants that may be an artifact. Genomic studies that fail to flag these regions are at risk of reporting incorrect candidates for downstream studies. We propose that the community apply the mask files provided here, until a more refined reference genome becomes available.

## Acknowledgements

This study is supported by U01DA043098 and R01DK099034.

---

**Supplementary Table 1.** Consistence across the three recent versions of the rat reference genome. The tables show the cross-tabulation of the number of genotype calls (in unit of 1,000) between rn4 and rn6 (upper table), and between rn5 versus rn6 (lower). The overall concordance is 0.98 between rn5 and rn6, and 0.97 between rn4 and rn6.

|  |  | rn6 |  |  |  |
| --- | --- | --- | --- | --- | --- |
|  |  | 0/0 | 0/1 | 1/1 | ./. |
| rn4 | 0/0 | 37,196.7 | 594 | 140.1 | 152 |
|  | 0/1 | 48 | 839 | 133.3 | 10.6 |
|  | 1/1 | 5.9 | 303.2 | 24958.4 | 155.1 |
|  | ./. | 537.8 | 498.6 | 1112.4 | 1920.8 |
|  |  | 0/0 | 0/1 | 1/1 | ./. |
| rn5 | 0/0 | 13908.1 | 374.7 | 59.8 | 240.6 |
|  | 0/1 | 48.2 | 1410.5 | 74.6 | 19.7 |
|  | 1/1 | 2.5 | 170.9 | 9606.8 | 351.7 |
|  | ./. | 72.3 | 65.6 | 103.3 | 364 |

**Supplementary Table 2.** Consistency of high-heterozygosity regions among three alignment methods. Shown are the total length of high-het regions for the 8 lines and with the use of the three aligners: *BWA, Bowtie2 and BWA-Stampy*. As the background level of heterozygosity differ among the three call sets, we applied different het fraction thresholds to define high-het windows: 0.25 for *BWA*, 0.175 for *Bowtie2*, and 0.2 for *Stampy*. The last three columns list concordance rates between heterozygous segments called by a pair of aligners, defined as intersect/union of the segments.

|  | *BWA* | *Bowtie* | *Stampy* | *BWA: Bowtie* | *BWA: Stampy* | *Bowtie: Stampy* |
| --- | --- | --- | --- | --- | --- | --- |
| **ACI** | 238,523,232 | 260,711,962 | 302,820,837 | 0.57 | 0.54 | 0.51 |
| **BN** | 171,689,865 | 189,496,386 | 213,984,578 | 0.61 | 0.60 | 0.58 |
| **BUF** | 223,339,767 | 247,060,856 | 289,737,173 | 0.55 | 0.55 | 0.49 |
| **F344** | 236,379,041 | 252,011,886 | 293,848,725 | 0.56 | 0.54 | 0.51 |
| **M520** | 243,992,401 | 243,643,168 | 294,212,048 | 0.58 | 0.54 | 0.5 |
| **MR** | 225,952,435 | 263,629,202 | 276,523,184 | 0.55 | 0.55 | 0.5 |
| **WKY** | 242,849,594 | 239,016,453 | 288,897,012 | 0.57 | 0.54 | 0.52 |
| **WN** | 228,435,487 | 249,395,735 | 298,824,484 | 0.53 | 0.52 | 0.47 |

**Supplementary Table 3.**
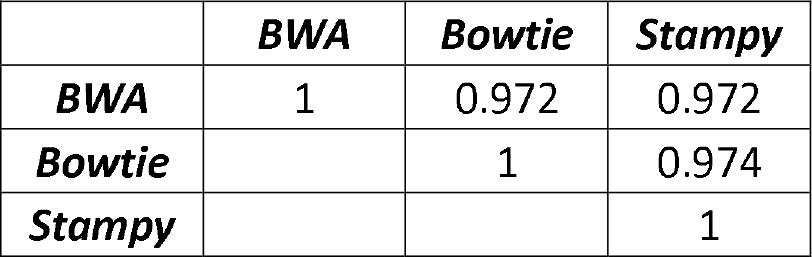
High levels of genotype concordance among variant calls obtained from different aligners (*BWA, Bowtie, BWA-Stampy*) followed by calling with UnifiedGenotyper. As before, we define concordance as the number of variant sites with the same genotype calls in both call sets divided by the number of variant sites with non-missing calls in both call sets.

|  | *BWA* | *Bowtie* | *Stampy* |
| --- | --- | --- | --- |
| *BWA* | 1 | 0.972 | 0.972 |
| *Bowtie* |  | 1 | 0.974 |
| *Stampy* |  |  | 1 |

**Supplementary Table 4.** Genotype counts and heterozygote frequencies in the eight lines. BN represents the reference genome and shows an outlier pattern for most metrics. 0/0, 0/1, 1/1 refers to Ref/Ref, Ref/Alt, and Alt/Alt genotypes. ./. refers to no-call due to low genotype quality.

| Strain | 0/0 | 0/1 | 1/1 | ./ | Het frequencies | Sites with > 2 alleles |
| --- | --- | --- | --- | --- | --- | --- |
| ACI | 6,249,928 | 2,096,993 | 5,421,143 | 2,385,991 | 0.15 | 251,129 |
| BN | 12,324,354 | 1,560,708 | 803,800 | 1,623,744 | 0.106 | 92,578 |
| BUF | 6,688,095 | 2,058,214 | 5,192,072 | 2,218,374 | 0.145 | 248,429 |
| F344 | 6,587,382 | 2,101,058 | 5,282,082 | 2,181,114 | 0.148 | 253,548 |
| M520 | 6,628,579 | 2,160,032 | 5,329,841 | 2,031,084 | 0.15 | 255,648 |
| MR | 6,736,380 | 2,111,216 | 5,243,452 | 2,063,655 | 0.147 | 250,481 |
| WKY | 6,599,091 | 1,976,801 | 5,130,247 | 2,456,145 | 0.142 | 242,900 |
| WN | 5,993,693 | 2,114,990 | 5,734,351 | 2,304,637 | 0.15 | 257,513 |

**Supplementary Table 5.**
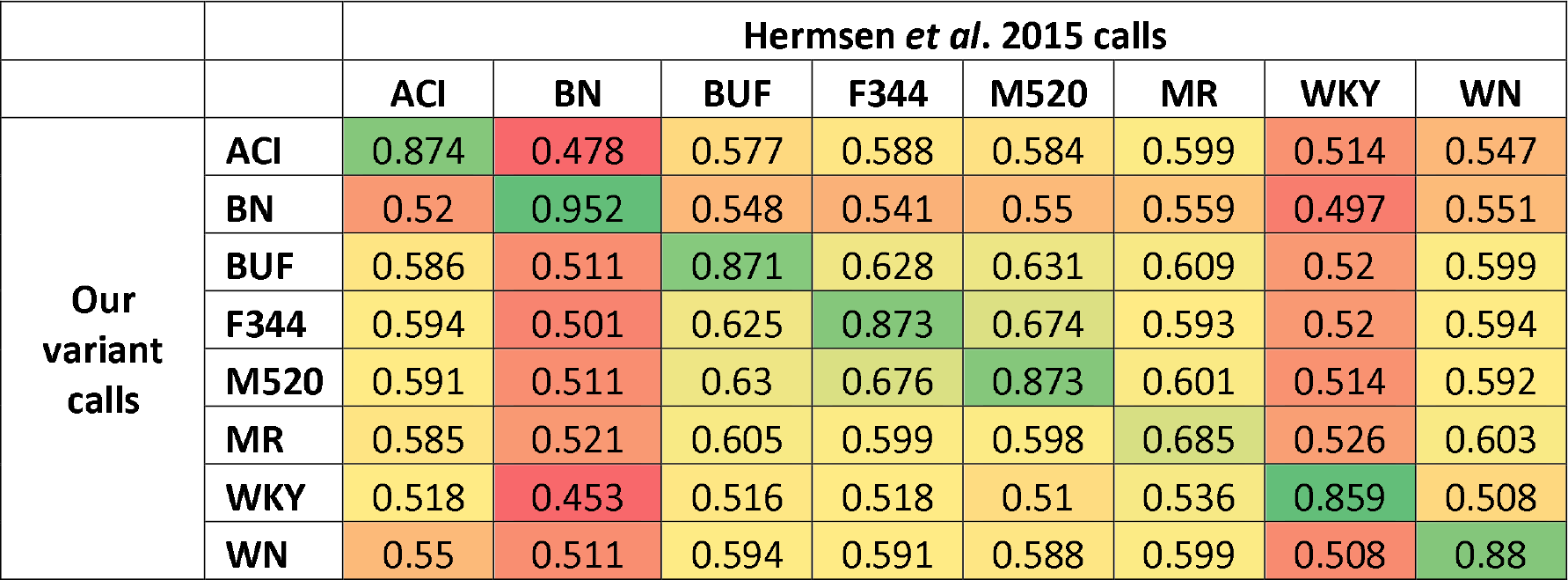
Concordance rate between our variant calls and the previous variant calls from Hermsen et al, 2015. The 8-by-8 table contains concordance values for all pair of samples. Here, we define concordance as the number of variant sites with the same genotype calls in both call sets divided by the number of variant sites with non-missing calls in both call sets.

**Supplementary Figure 1.**
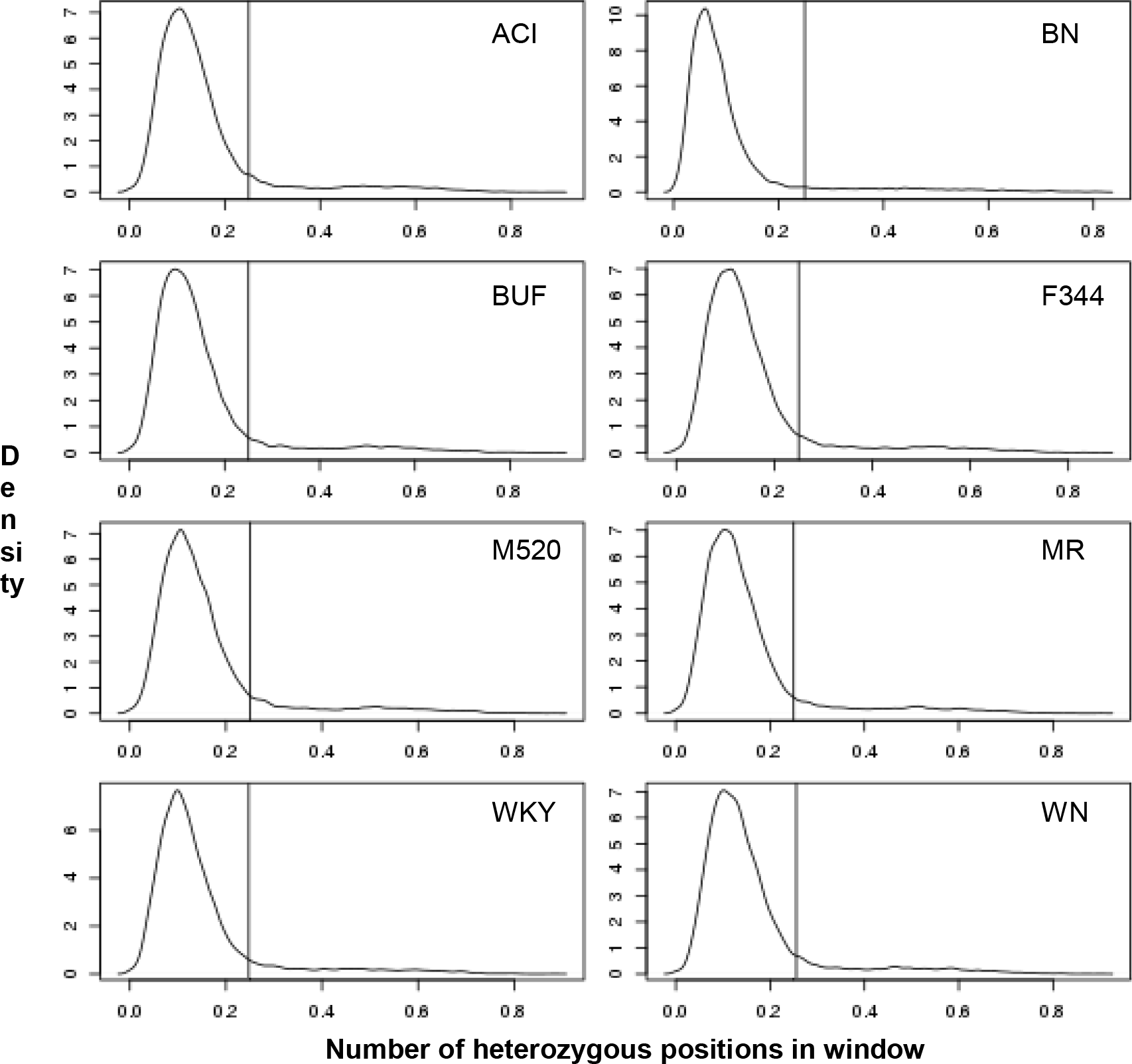
Distribution of the heterozygosity level in 1000-SNV windows for each of the eight lines, suggesting that 0.25 is a reasonable cutoff for defining high-het windows.

**Supplementary Figure 2.**
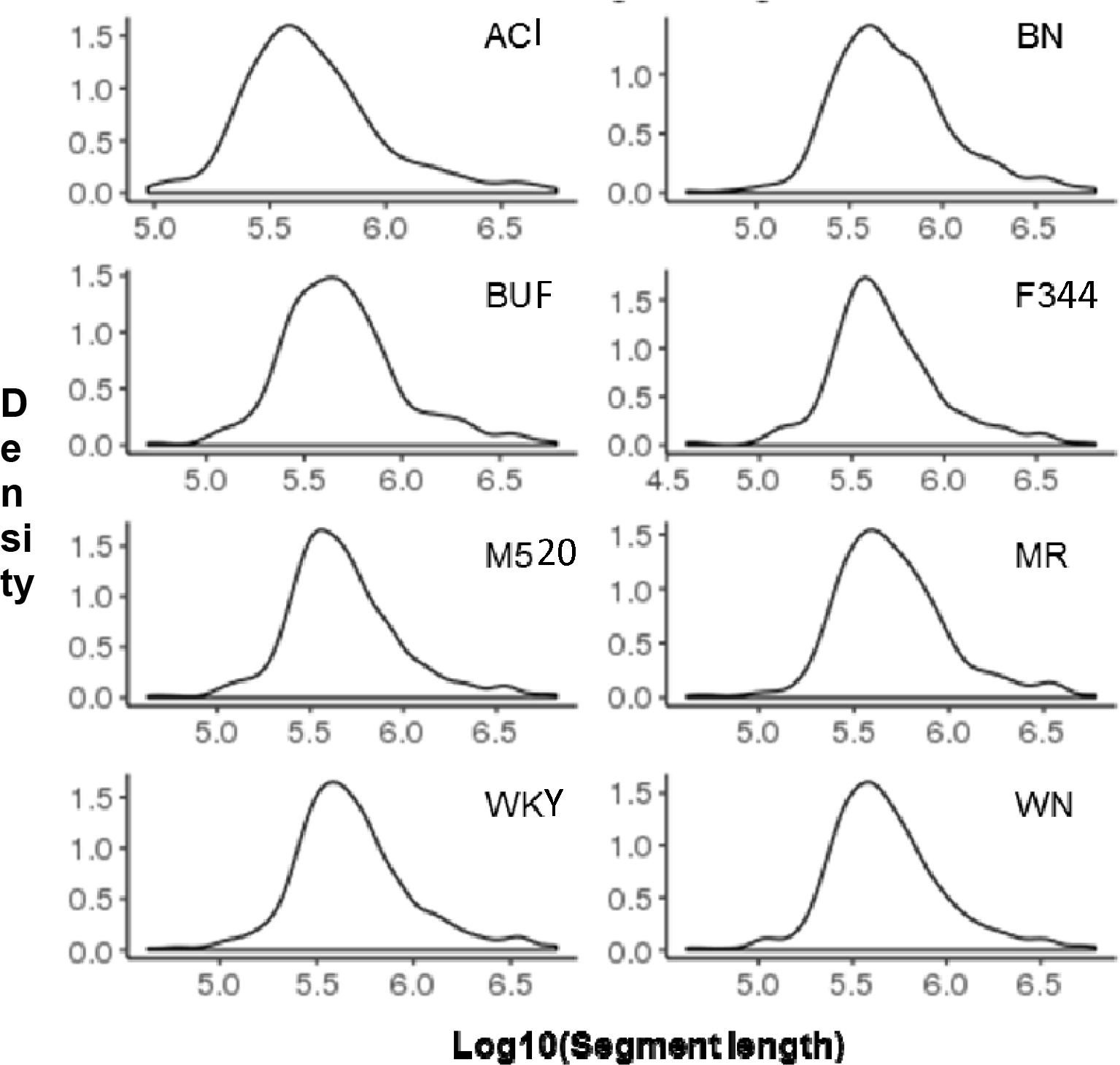
Distribution of high-het segment lengths in the 8 lines.

**Supplementary Figure 3.**
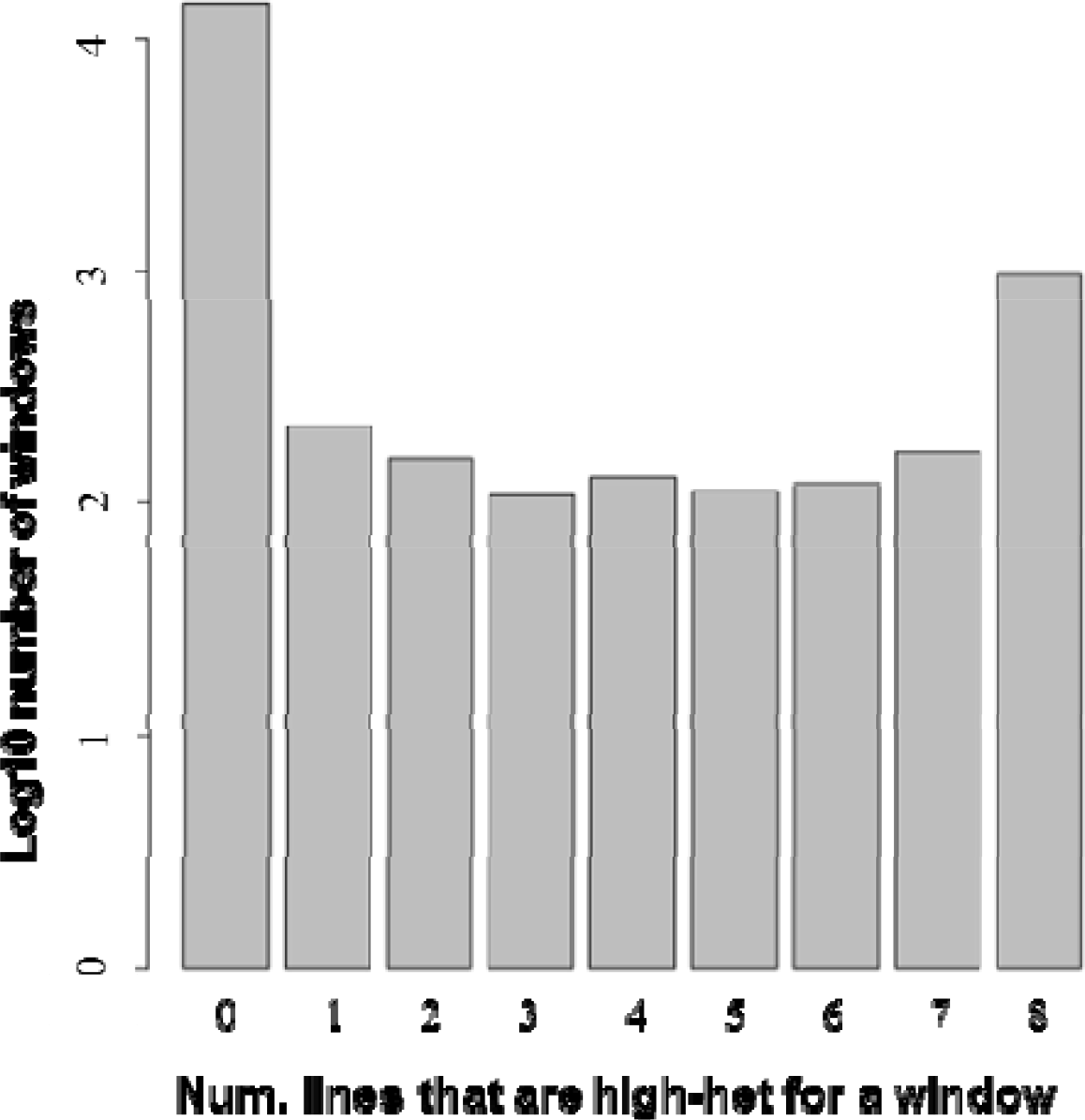
Histogram of the number of 1000-SNV windows that are high-het (heterozygosity > 0.25) in, from left to right, 0, 1, …, 8 founder lines.

**Supplementary Figure 4.**
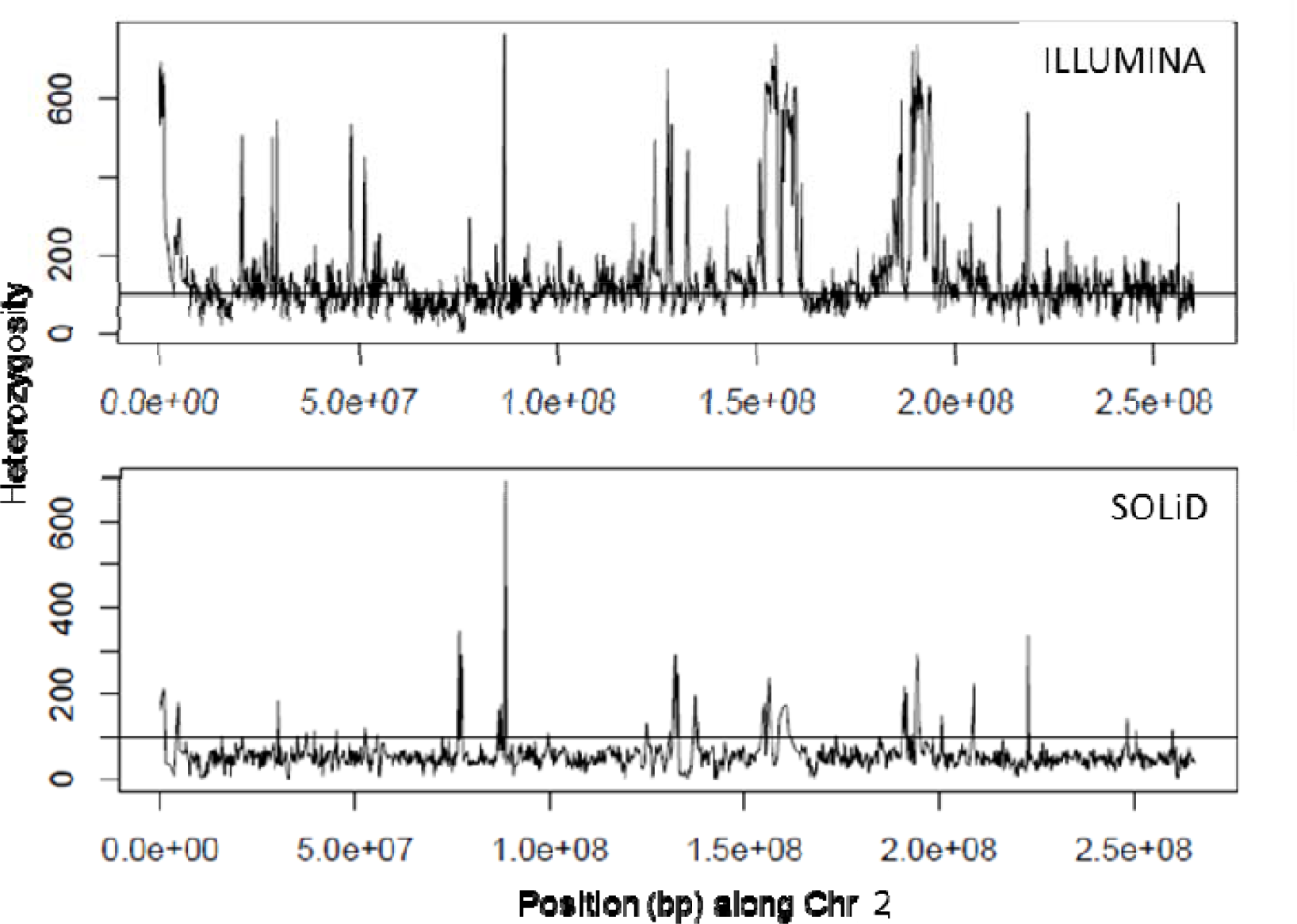
Similar regional patterns of high-het segments in Chr2 between this study and the previously reported SOLiD calls. The x-axis is displayed as base positions of the 1000-SNV windows rather than the window IDs, because the two call sets (Illumina and Solid) contained different numbers of variant sites, and the 1000-SNV windows are mismatched between the two. The horizontal line represents heterozygosity of 100.

**Supplementary Figure 5.**
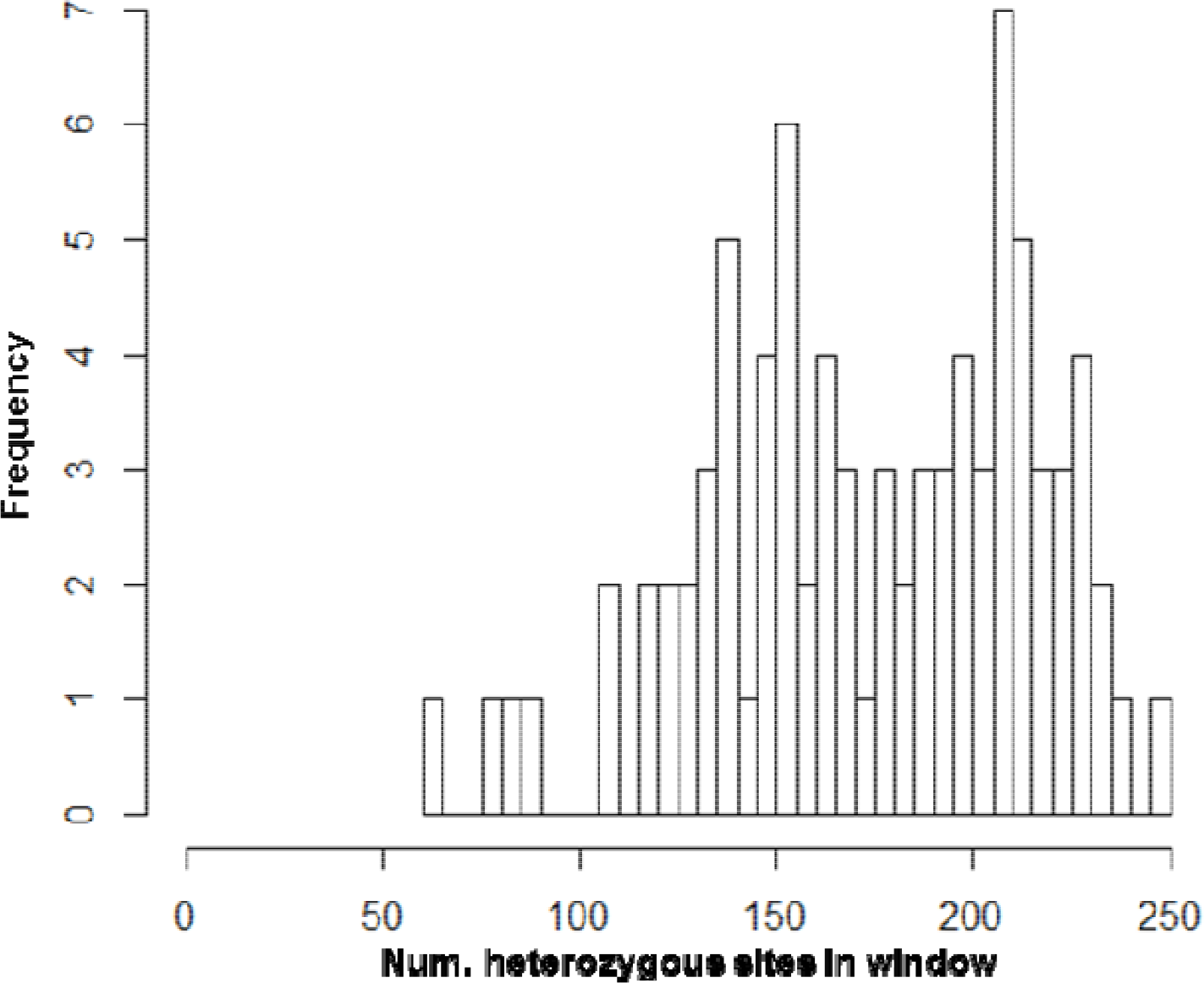
Distribution of heterozygosity rates in windows that are initially classified as low-het, but are flanked by high-het windows on both sides. The bimodal distribution suggests those with >175 hets may be merged with flanking high-het segments.

